# Disrupted Integration–Segregation Balance in the Intact Hemisphere in Chronic Spatial Neglect

**DOI:** 10.1101/2025.10.20.683071

**Authors:** Yusaku Takamura, Marine Lunven, Yvonne Serhan, Clémence Bourlon, Smadar Ovadia-Caro, Paolo Bartolomeo

## Abstract

Spatial neglect is a common and disabling consequence of right hemisphere stroke, characterized by a failure to attend to the contralesional left space, and frequently persists into the chronic stage. There is robust evidence on the role of right-hemisphere frontoparietal dysfunction, interhemispheric structural disconnection and maladaptive activity in the left hemisphere in the persistence of neglect. However, the specific impact of right frontoparietal dysfunction on the functional (re)organization of the left hemisphere remains poorly understood. In this study, we introduce a novel application of functional connectivity gradient analysis to investigate macroscale functional reorganization in the non-lesioned left hemisphere of patients with chronic left spatial neglect. Focusing on resting-state fMRI data, we demonstrate that abnormal segregation patterns in the left frontoparietal and default mode networks are robustly associated with neglect severity and spatial attentional bias. Notably, the principal gradient—typically capturing a global unimodal-to-transmodal hierarchy—was altered in these patients, suggesting a reorganization favoring lateralized unimodal networks. Single-subject analyses confirmed the presence of this pattern in 11 of the 13 patients included in the study. We also show that the structural integrity of the left inferior fronto-occipital fasciculus (IFOF) plays a key role in shaping these functional dynamics. These findings reveal a previously overlooked aspect of neglect pathophysiology: the maladaptive dominance of the non-lesioned hemisphere’s intrinsic architecture. By combining innovative gradient-based metrics with classical lesion approaches, our study offers a new framework for understanding neglect as an emergent property of large-scale network imbalance, with clinical implications for diagnosis and intervention, and theoretical consequences for models of hemispheric asymmetries and conscious access.

## Introduction

Left spatial neglect is a disabling neuropsychological condition that frequently follows right hemisphere stroke, and is characterized by a failure to report, respond to, or orient toward stimuli presented on the contralesional side (Heilman et al., 2008). In the acute phase, up to 61% of right hemisphere stroke patients present signs of contralesional neglect, compared to only 22% of those with left hemisphere damage (Cazzoli et al., 2025). Furthermore, neglect after right-hemisphere lesions is typically more severe and persistent, with about 40% of patients exhibiting chronic symptoms months after their stroke (Overman et al., 2024). These deficits profoundly impact daily living, rehabilitation outcomes, and quality of life (Bosma et al., 2020; Moore et al., 2021; Oh-Park et al., 2014). Neglect is widely regarded as an attentional disorder arising from disruptions in large-scale brain networks rather than from isolated cortical lesions (Bartolomeo et al., 2007; Corbetta & Shulman, 2011; Doricchi et al., 2008). Structurally, disconnections within frontoparietal networks—particularly due to damage to the superior longitudinal fasciculus (SLF)—coupled with disruption of interhemispheric callosal fibers, contribute to the onset and chronic persistence of neglect (Kaufmann et al., 2024; Lunven et al., 2015, 2019; Thiebaut de Schotten et al., 2014). Specifically, exploration of peripersonal space and awareness of left-sided events (Martín-Signes et al., 2024) are disrupted by damage to the ventral (Urbanski et al., 2011) and intermediate (Thiebaut de Schotten et al., 2005, 2008) branches of the right-hemisphere SLF, which connect the ventral attention network (VAN) and the dorsal attention network (DAN), respectively (Corbetta & Shulman, 2011; de Schotten et al., 2011). Impaired interhemispheric coordination between the right and left DANs also contributes to neglect signs (Corbetta & Shulman, 2011; He et al., 2007). Additional contributions arise from dysfunction of the default mode network (DMN), whose hyperactivity, loss of inhibition, and antagonistic interaction with the DAN are implicated in the pathophysiology and recovery of neglect (Bonnelle et al., 2011; Fox et al., 2005; Golland et al., 2007; Raichle et al., 2001; Ramsey et al., 2016; Weissman et al., 2006).

While the lesioned right hemisphere has been the primary focus of much research, emerging evidence indicates that the intact left hemisphere may also undergo functional reorganization in patients with chronic neglect. This reorganization may support compensatory processes or, in some cases, reflect maladaptive changes, but the mechanisms and functional consequences remain incompletely understood (Bartolomeo, 2019; Di Pino et al., 2014, Lunven et al 2015, Corbetta & Shulman 2011). Supporting this complex picture, activity in the left frontal cortex has been reported to precede left-sided omissions in chronic neglect (Rastelli et al., 2013), whereas interventions such as inhibitory neuromodulation (Oliveri et al., 1999) or, in a case report, a second stroke affecting the left prefrontal cortex (Vuilleumier et al., 1996), have been associated with improved performance. Recovery has also been linked to normalization of interhemispheric connectivity, increased network segregation, and altered interactions between the DMN and attention networks across both hemispheres (Corbetta & Shulman, 2011; He et al., 2007; Ramsey et al., 2016; Umarova et al., 2016). Together, these findings highlight a complex interplay between compensatory and maladaptive plasticity, as well as substantial inter-individual variability, representing an important knowledge gap in understanding the evolution of neglect (Bartolomeo, 2019; Di Pino et al., 2014).

Here, we address this gap by introducing a novel approach based on functional connectivity gradients (FGs) in the non-lesioned left hemisphere of patients with chronic left neglect. Unlike traditional connectivity analyses, which focus on discrete regions or pairwise connections and may overlook the brain’s global or hierarchical organization, FGs offer a low-dimensional, continuous representation of whole-brain architecture. FGs capture the spatial progression from unimodal to transmodal regions and reflect the underlying balance between functional segregation and integration, thereby providing a powerful new lens for investigating large-scale reorganization in neglect (Margulies et al., 2016; Serhan et al., 2025). This approach has proven highly effective in detecting alterations across neurological and psychiatric disorders (Dong et al., 2023; Hong et al., 2019; Rosa et al., 2024; Tan et al., 2023), including stroke (Bayrak et al., 2019). Consequently, FGs represent a promising framework for investigating distributed, macroscale brain network reorganizations related to attentional processes and their disruption in neglect chronicity (Ramsey et al., 2016; Siegel et al., 2018).

In the present study, we apply functional connectivity gradient analyses to describe large-scale network organization in the non-lesioned left hemisphere of patients with chronic left neglect. Our approach is exploratory, examining how individual differences in gradient architecture relate to behavioral measures. By focusing on the chronic phase (>3 months post-stroke), we aim to characterize stable, long-term patterns of functional organization relatively free from acute confounds such as edema or transient diaschisis (Carrera & Tononi, 2014; Grefkes & Fink, 2020; Ramsey et al., 2016; Rehme et al., 2011; Siegel et al., 2018). This work provides a basis for future studies investigating how variability in the intact hemisphere may relate to recovery or persistent deficits in larger and more diverse stroke cohorts.

## Methods

### Participants

The data analyzed in this study were previously collected and published by Lunven et al., (2019) in a study devoted to the effects of prism adaptation in chronic neglect (Lunven et al., 2019). The study received ethical approval from Inserm (C10-48) and authorization from the Ile-de-France I Ethics Committee (2011-juin-12642). All participants provided written informed consent to participate in accordance with the Declaration of Helsinki.

Here we focus on the behavioral data acquired two hours before prism adaptation. Neuroimaging data were collected before prism adaptation, depending on each participant’s availability and the original study protocol (see details in Lunven et al., (2019)). The dataset includes 13 patients with chronic visual neglect (Age: 63.92 ± 9.79 years, education level: 12.20 ± 2.53 years of formal education) following a right hemisphere stroke (more than 90 days post-stroke) and 10 neurologically healthy control (HC) participants matched for age and education (age: 63.4 ± 4.24 years, education level: 11.76 ± 2.99 years of formal education). Detailed demographic, clinical, and neuropsychological characteristics are presented in Table 1. Neuropsychological assessments included standardized tests commonly used to evaluate neglect sign: letter cancellation (Mesulam 2000), copy of a linear drawing of a landscape (Gainotti et al., 1989), drawing from memory of a daisy (Rode et al., 2001), text reading (Azouvi et al., 2006), bisection of five 20-cm horizontal lines (Azouvi et al., 2006; Urbanski & Bartolomeo, 2008) and the landmark task (Harvey & Milner, 1999). For each test, individual performance was converted into a standardized score, expressed as a percentage of maximum possible performance, with higher values indicating greater neglect severity. These standardized scores were then combined to create a composite neglect score, providing a global measure of spatial neglect across multiple modalities and task demands. This approach captures both perceptual and motor components of neglect, as well as deficits in exploratory behavior and representational space. Standard scoring procedures were applied as described in Lunven et al., (2019). Lesion overlap maps for all patients and analysis pipelines are presented in Figure 1.

**Figure 1.**
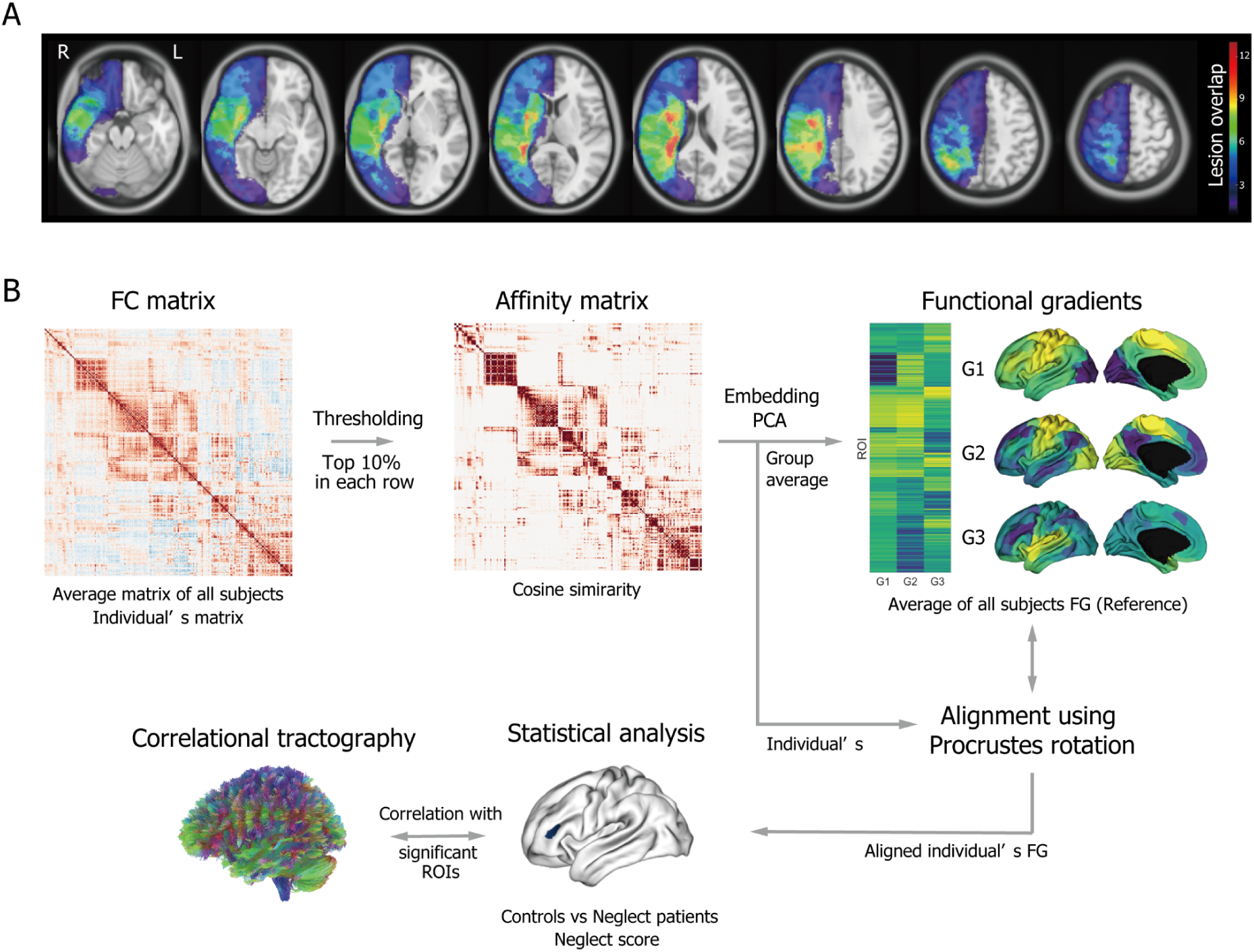
Lesion overlap map and schematic of the analysis pipeline. (A) Lesion overlap map for the 13 patients included in the study. Warmer colours indicate higher overlap across individuals. (B) Overview of the analysis pipeline. Functional connectivity (FC) matrices were derived from preprocessed resting-state fMRI time series at both the group-level and individual-level. An affinity matrix was constructed by using pairwise cosine similarity for the thresholded FC matrix. Functional connectivity gradients were then extracted using principal component analysis (PCA) and projected into a low-dimensional embedding. Individual gradient maps were aligned to a group-averaged template using Procrustes alignment. Statistical comparisons were performed and regions showing significant differences were subsequently examined as the dependent variables to be explained by structural connectivity.

**Table 1.** Demographic and clinical characteristics of patients.

|  | **Cutoff score** | **Neglect (n = 13)**<br>**Mean (SD)** |
| --- | --- | --- |
| Age (years) | N/A | 63.92 (9.79) |
| Weeks since stroke | N/A | 82.68 (63.27) |
| Lesion volume (cm3) | N/A | 200.41 (147.41) |
| MMSE (pts) | < 23 | 27 (1.92) |
| Neglect score (%) | N/A | 31.75 (22.28) |
| Letter cancellation total hits<br>(max = 60) | N/A | 47.53 (12.85) |
| Letter cancellation L-R difference in omissions<br>(-30 to +30) | >3 | 6.92 (5.79) |
| Gainotti drawing<br>(max = 6) | <6 | 4.53 (1) |
| Reading<br>(max = 116) | <116 | 103.15 (24.15) |
| Landmark<br>“Left shorter responses” | >8 | 7.69 (3.24) |
| Line bisection<br>(deviation mm) | >6.5 | 19.05 (24.27) |
| Daisy drawing<br>(max = 3) | <3 | 1.76 (0.89) |

MMSE: Mini Mental State Examination

### Data acquisition

Patients and controls underwent MRI on a Siemens 3 T VERIO TIM system equipped with a 32-channel head coil, generating T1-weighted structural images (MPRAGE), resting-state blood oxygen level-dependent (BOLD) functional scans (rsfMRI), and diffusion-weighted images (DWI). The details of T1-weighted images, lesion mask segmentations, and DWI have been described in a previous study (Lunven et al., 2019). Functional images were acquired with repetition time = 2.99s, echo time = 26 ms, flip angle = 90°, slice thickness of 3 mm, and 200 repetition times (scanning duration of approximately 10 min).

### fMRI preprocessing

The data were transformed from DICOM to BIDS format using HeuDiConv (Halchenko et al., 2024). Next, lesion masks were included in each BIDS-style folder to reduce the warping of healthy tissue into damaged areas and vice versa during spatial normalization (Brett et al., 2001). The following description regarding the preprocessing pipeline was automatically generated by fMRIPrep. It is supported under the CC0 license. Results included in this manuscript come from preprocessing performed using fMRIPrep 24.1.1 (Esteban et al., 2019), which is based on Nipype 1.8.6 (Gorgolewski et al., 2011).

### Anatomical data preprocessing

Anatomical T1-weighted images were preprocessed using standard pipelines on fMRIprep, including intensity non-uniformity correction (N4BiasFieldCorrection (Tustison et al., 2010)), skull-stripping (ANTs (Avants et al., 2008)), and tissue segmentation into gray matter, white matter, and cerebrospinal fluid using FSL’s FAST (Zhang et al., 2001). Cortical surface reconstruction was performed with FreeSurfer’s ‘recon-all’ (Dale et al., 1999), and segmentation outputs were reconciled using Mindboggle tools (Klein et al., 2017). Spatial normalization to MNI152NLin2009cAsym standard spaces was achieved via nonlinear registration with ANTs. Grayordinate resampling to fsLR was performed using Connectome Workbench tools (Glasser et al., 2013). Following the fMRIPrep recommendations (https://fmriprep.org/en/latest/workflows.html), lesion masks were included to optimize spatial normalization and minimize distortion of damaged tissue (Brett et al., 2001).

### Functional data preprocessing

Functional images were preprocessed using standard pipelines. Head motion correction was performed using FSL’s mcflirt (Jenkinson et al., 2002), with motion parameters estimated relative to a reference volume generated by fMRIPrep. The BOLD reference was co-registered to the T1w image using boundary-based registration (bbregister (Greve & Fischl, 2009)), using six degrees of freedom. Confound time series were computed, including framewise displacement (Jenkinson et al., 2002; Power et al., 2014), DVARS, and global signals from white matter, cerebrospinal fluid, and the whole brain. BOLD time series were mapped to the fsLR surface space using the Connectome Workbench (Glasser et al., 2013), generating CIFTI-91k grayordinate files. All spatial transformations (motion correction, co-registration, normalization) were composed into a single resampling step using cubic B-spline interpolation. Lesion masks were incorporated during spatial normalization to reduce distortion associated with damaged tissue (Brett et al., 2001) following to the fMRIprep recommendations (https://fmriprep.org/en/latest/workflows.html). Subsequent preprocessing pipelines were relied fMRIPrep and Nilearn v0.10.4 (Abraham et al., 2014). This process included: (i) removal of the first 3 volumes to mitigate signal equilibration effects; (ii) exclusion of time points with excessive motion (>2 mm or >0.02 radians) or global signal outliers (>3 SD (Demertzi et al., 2019)); (iii) nuisance regression of motion parameters, white matter, cerebrospinal fluid, and global signal (Fox et al., 2005; Wang et al., 2024); and (iv) band-pass filtering (0.01–0.1 Hz (Bayrak et al., 2019)). Global signal regression was included to improve brain–behavior associations (Li et al., 2019).

### Functional connectivity gradients and statistical analysis

The analysis workflow is summarized in Figure 1B. Functional connectivity (FC) matrices were constructed using the Schaefer atlas (Schaefer et al., 2018), comprising 400 cortical parcels across 7 networks, plus 54 subcortical regions (Tian et al., 2020), for a total of 454 region of interest (ROI) Spatial smoothing was not applied, as signal averaging within ROIs effectively reduces high-frequency noise (Alakörkkö et al., 2017). FC matrices (454 × 454) were computed using Pearson’s correlation coefficient. To avoid confounds from large lesions in patients, 227 right-hemisphere ROIs were excluded, consistent with previous MEG research (Rastelli et al., 2013).

FC gradients were computed using Brainspace (Vos de Wael et al., 2020). The connectivity matrix was thresholded to retain the top 90% of correlation coefficients per row, and pairwise cosine similarities was calculated (Bayrak et al., 2019). Principal component analysis (PCA) was used for embedding, as it offers robust reliability and predictive validity for gradient mapping (Hong et al., 2020). Based on prior research, we considered selecting the top three gradients to ensure a balanced representation of connectivity patterns (Bayrak et al., 2019). However, to account for the restriction of our analysis to the left hemisphere only, we included up to 20 gradients. Additional components were not included when the cumulative variance explained by the first three gradients reached the levels reported in previous stroke studies (50.84% (Bayrak et al., 2019) and 41.63% (Koba et al., 2025)). Individual gradients were aligned to the group average using procrustes rotation (90,93,94). To compute the distance of distribution between the two groups, we also calculated the Wasserstein distance (Ramdas et al., 2017), after transforming the group-level gradients into z-scores, which quantifies the cost of transforming one distribution into another.

Statistical analysis was performed with Brainstat (Larivière et al., 2023) using linear regression models and followed by multiple comparison correction with false discovery rate (FDR) method (Benjamini & Hochberg, 1995). Lesion volume and visual field deficits were included as nuisance variables (Lunven et al., 2019). We focused on the line bisection and letter cancellation tasks, in addition to the global neglect score, to capture complementary aspects of left spatial neglect. Line bisection reflects perceptual and representational spatial bias, while letter cancellation is sensitive to deficits in exploratory attention (Azouvi et al., 2006; Mesulam, 2000; Urbanski & Bartolomeo, 2008). Together, these measures represent different neglect subtypes (Toba et al., 2018; Verdon et al., 2010) and maintain consistency with the composite score (Lunven et al., 2015, 2019). Despite occasional reports of low sensitivity (notably when lines are right-aligned on the (Ferber & Karnath, 2001), 20-cm line bisection remains clinically informative: in Azouvi et al. (2023), 20-cm line bisection such as the one used here showed meaningful discrimination and was retained in the most parsimonious high-specificity screening model. Using these measures, we compared controls and neglect patients and examined associations between functional connectivity gradients and neglect severity. Two outlier scores exceeding two standard deviations were excluded from line bisection and global neglect measures, the excluded patients’ scores were 76.03% and 81.11% for Neglect severity, and 73 mm and 70 mm for the line bisection test. There were no excluded patients in the letter cancellation test.

The analysis pipeline incorporated publicly available scripts and functions, including hcp-utils (https://github.com/rmldj/hcp-utils/tree/master), stroke_preprop (Bayrak et al., 2019) https://github.com/sheyma/stroke_preprop, and autism_gradient_asymm (Wan et al., 2023) https://github.com/wanb-psych/autism_gradient_asymm/tree/master. Finally, we computed partial r-squared value and Cohen’s F as measures of effect size (Cohen, 2013; Gravetter et al., 2021; Selya et al., 2012).

### Diffusion MRI preprocessing

The preprocessing pipeline has been described in a previous study (Lunven et al., 2019). As a preprocessing step, brain extraction was performed using BET as implemented in FSL (Smith et al., 2004). Diffusion-weighted datasets were simultaneously corrected for subject motion and eddy current-induced geometric distortions using ExploreDTI (http://www.exploredti.com) (Leemans & Jones, 2009). Subsequently, the diffusion tensor model was fitted to the data using Levenberg-Marquardt nonlinear regression (Marquardt, 1963). Subsequently, preprocessed diffusion data were processed in DSI-Studio (“Hou” version: https://dsi-studio.labsolver.org/). The following descriptions were based on the boilerplate automatically generated by DSI-Studio. It is supported under the CC BY-NC-SA 4.0 license. A total of 23 diffusion MRI scans were included in the connectometry database with the 2.000 mm in-plane resolution. The diffusion data were reconstructed in the MNI space using q-space diffeomorphic reconstruction (Yeh & Tseng, 2011) to obtain the spin distribution function (Yeh et al., 2010). A diffusion sampling length ratio of 2 was used. The output resolution in diffeomorphic reconstruction was 1.72 mm isotropic. The restricted diffusion was quantified using restricted diffusion imaging (Yeh et al., 2017). The tensor metrics were calculated using DWI with b-value lower than 1750 s/mm². The quantitative anisotropy (QA) was extracted as the local connectome fingerprint and used in the connectometry analysis.

### Correlational tractography to explain the altered functional gradients

To understand the structural connectivity that underpins the significantly altered functional gradients, we used correlational tractography on DSIstudio (Kaufmann et al., 2024). DSIstudio automatically generated the following descriptions. Correlational tractography (Yeh et al., 2021) was derived to visualize pathways that have QA correlated with significant ROIs. A nonparametric Spearman partial correlation was used to derive the correlation, and the effect of lesion volume and presence of visual field deficits was removed using a multiple regression model. The statistical significance of the correlation was examined using a permutation test (Yeh et al., 2016). A total of 23 subjects were included in the analysis. A T-score threshold of 3.5 was assigned in the fiber tracking algorithm (Yeh et al., 2013). Cerebellum was excluded. The lesion region was removed from the tracking. A seeding region was placed in the non-lesioned cerebral regions. The tracks were filtered by topology-informed pruning (Yeh et al., 2019) with 16 iterations. An FDR threshold of 0.05 was applied, with 4000 randomized permutations of group labels used to estimate the null distribution of track length. Significant results were automatically segmented on DSIstudio to underlying fiber tracks.

### Individual analyses of functional gradients

Neglect is often considered a heterogeneous condition, raising the possibility that patients may exhibit different patterns of resting-state brain activity. To assess potential individual differences, we conducted single-case statistical analyses comparing each neglect patient with the HC group (Crawford et al., 2009, 2010; Cubelli & Della Sala, 2017). Singcar package (Rittmo & McIntosh, 2021) was used to perform the Crawford test using the Bayesian test of deficit with 100,000 iterations on R (Version 4.2.2). Statistical significance, indicating whether the neglect patients followed the group average, was assessed by testing whether the 95% confidence interval of the effect size excluded zero.

## Results

### Spatial patterns of functional connectivity gradients

Figure 2A displays the first three FC gradients for HC and neglect patients. Together, these gradients accounted for 58.02% of the variance (23.41%, 21.11%, and 13.49%, for Gradients 1-3, respectively), with broadly similar spatial distributions across groups. Figure 2B shows pairwise scatter plots illustrating the low-dimensional embedding of cortical connectivity. Gradient 1 spanned visual to somatomotor networks, Gradient 2 extended from transmodal regions, i.e., the DMN, to unimodal sensory networks, and Gradient 3 separated frontoparietal control network (FPCN) and DAN, from VAN and perisylvian somatomotor networks. The general arrangement of functional gradients was preserved in neglect patients, although Gradient 3 showed greater differences than Gradients 1 and 2 as indicated by the Wasserstein distance (Gradient 1: 0.042; Gradient 2: 0.065; Gradient 3: 0.142). In particular, subtle differences were observed in the dispersion of specific functional networks.

**Figure 2.**
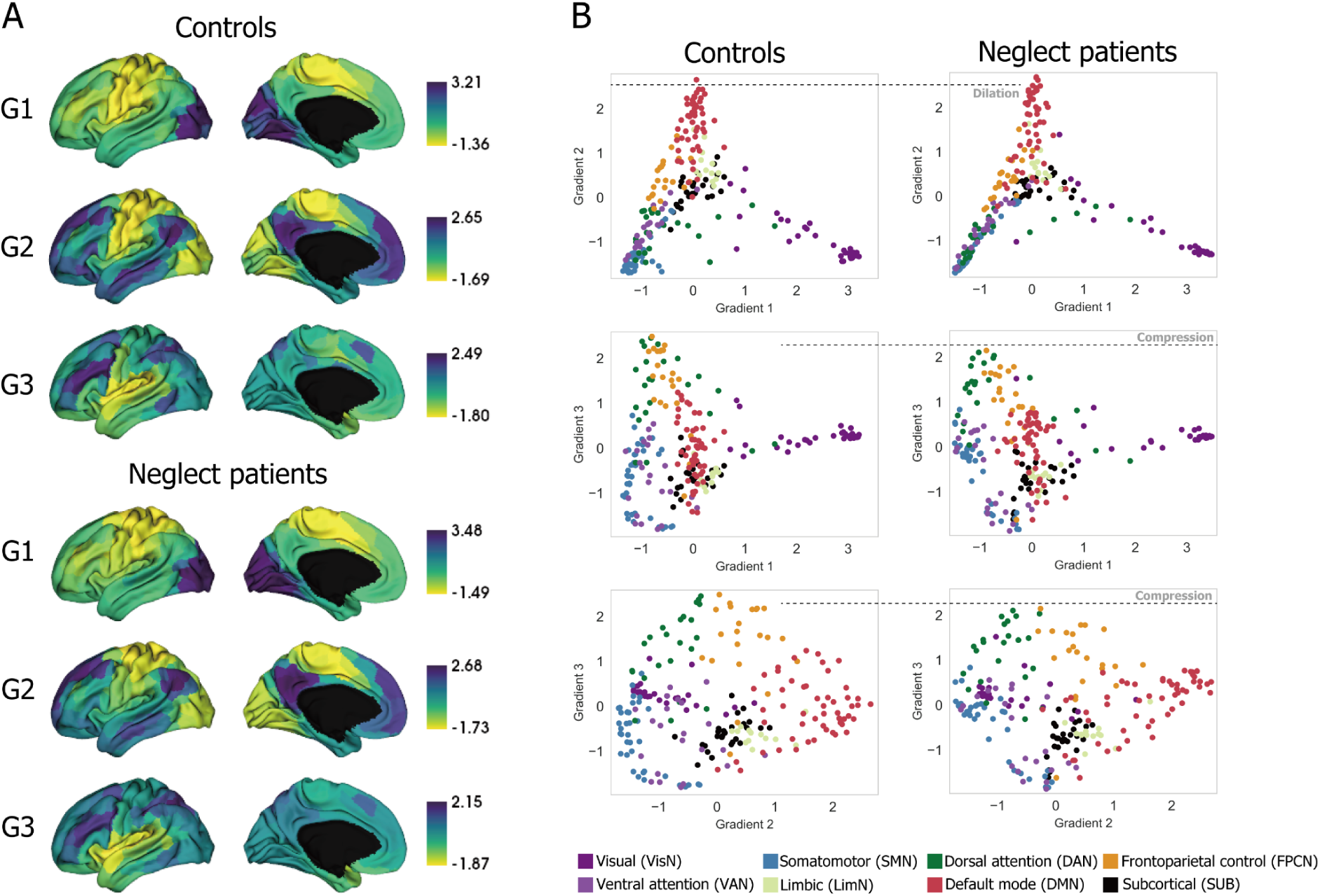
The general arrangement of functional networks along gradients is preserved in the left hemisphere of neglect patients and subtle differences are expressed in altered dispersion. (A) Functional connectivity gradients derived from resting-state fMRI in neglect patients and healthy controls. The top three functional gradients are shown. Gradient 1 primarily distinguishes between unimodal networks, spanning an axis from visual to somatomotor regions. Gradient 2 extends from transmodal to unimodal networks, with one extreme anchored in the default mode network and the other in sensory networks. Gradient 3 reflects task-related networks, delineating an axis from the frontoparietal control and dorsal attention networks to the ventral attention and somatomotor perisylvian networks. (B) Pairwise scatter plots of the first three gradients highlight group-level differences, with the most prominent divergence observed in Gradient 3. Compared to healthy controls (HC), patients with neglect show reduced dispersion in regions associated with the frontoparietal control and dorsal attention networks.

Compared to HC, neglect patients showed reduced dispersion within the FPCN and DAN along Gradient 3 and increased dispersion in the DMN along Gradient 2.

### Relationships between functional connectivity gradients and neglect

Figure 3 summarizes the statistical analyses of FC gradients in relation to neglect. The whole-brain comparison of gradient distributions between neglect patients and HC showed significant differences in the lateral prefrontal cortex, within the frontoparietal control network in Gradient 3 (Cont_PFC1_2: t = –4.858; pFDR = 0.024; partial R^2^ = 0.554; Cohen’s F = 1.114, Figure 3A). Neglect patients exhibited gradient values shifted closer to zero in this region compared to controls (t-test: t = –3.592; p = 0.001; Cohen’s d = –1.421; Figure 3B). In neglect patients, correlations between gradient values and line bisection performance revealed a negative association in the frontoparietal control network and a positive association in DMN regions (Figure 3C, Cont_PFC1_6: t = -5.874; pFDR = 0.028; partial R^2^ = 0.831; Cohen’s F = 2.220; Default_Temp_4: t = 5.500; pFDR = 0.034; partial R^2^ = 0.812; Cohen’s F = 2.079; Default_Par_4: t = 7.562; pFDR = 0.015; partial R^2^ = 0.890; Cohen’s F = 2.858; Default_Par_6: t = 8.324; pFDR = 0.015; partial R^2^ = 0.908; Cohen’s F = 3.146; Default_PFC_11: t = 6.080; pFDR = 0.028; partial R^2^ = 0.841; Cohen’s F = 2.298; Default_pCunPCC_7: t = 5.953; pFDR = 0.028; partial R^2^ = 0.835; Cohen’s F = 2.250). The regression analysis using extracted regional gradient values, adjusted for nuisance covariates, confirmed these associations (Figure 3D; Cont_PFCl_6: t = –5.874; p = 0.001; R² = 0.841; DMN mean: t = 11.026; p < 0.001; R² = 0.931). We further examined correlations between gradient values and left–right differences in the letter cancellation test and neglect severity, but none of these relationships reached statistical significance. Finally, violin plots summarized the distribution of network-level gradients (Figure 3E), highlighting a group difference in the frontoparietal control network for Gradient 3 (t = 3.845; pFDR = 0.003; Cohen’s D = 1.618), alongside increased dispersion in DMN in Gradient 2 in neglect patients.

**Figure 3.**
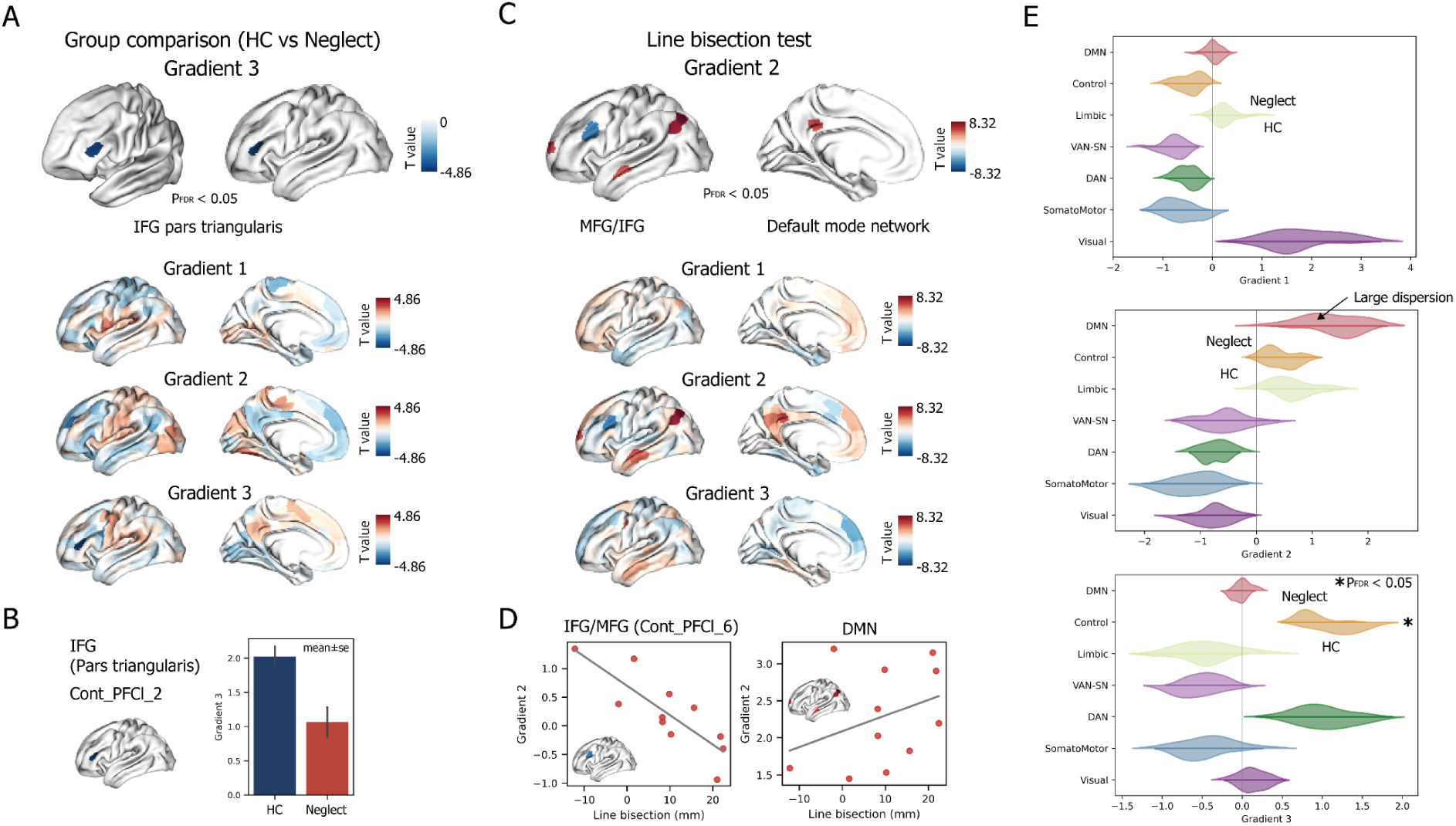
Areas of the frontoparietal network show reduced integration while areas of DMN show enhanced integration in neglect patients. (A) T-statistics across the three gradients comparing healthy controls (HC) and neglect patients, with statistically significant regions highlighted in the top row. A significant group difference was observed in the lateral prefrontal cortex, attributed to the frontoparietal control network. This region anatomically corresponds to the pars triangularis of the inferior frontal gyrus (IFG). (B) Bar plots for Cont_PFCl_2, which showed a significant difference between the HC and neglect groups, corresponding to panels (A) and (C). Bars represent group means, and error bars indicate the standard error of the mean (SEM). (C) T-statistics of the correlation between three gradients scores and line bisection test, with statistically significant regions highlighted in the top row. A significant negative correlation was observed in the lateral prefrontal cortex, attributed to the frontoparietal control network (Cont_PFCl_6). This region anatomically corresponds to the pars triangularis of the IFG and middle frontal gyrus (MFG). Gradient 2 score at DMN also showed significant correlation with line bisection score. (D) Correlations between neglect severity (deviation of line bisection) and gradient values in two representative regions. Left: A negative correlation was observed in the Cont_PFCl_6. Right: A positive correlation was observed in the DMN regions. Both scatter plots indicate that a greater rightward bias in line bisection was associated with gradient 2. (E) Violin plot showing the average of three gradient networks in neglect and HC.Significant difference was observed in the frontoparietal control network in gradient 3. Large dispersion values were also observed in the default-mode network in gradient 2 (depicted by arrow).

### Connectometry analysis explaining altered functional connectivity gradients

Connectometry analysis revealed a positive correlation between Gradient 3 values in the left prefrontal cortex (Cont_PFCl_2) and QA values in the anterior commissure (AC), and white matter pathways encompassing the left occipito-temporal region, such as the inferior fronto-occipital fasciculus (IFOF), and arcuate fasciculus (AF) (Figure 4A). We observed overlap between the AC and IFOF, consistent with previous reports that their dorsoventral distributions overlap (Baudo et al., 2022). A positive correlation was also observed between the mean Gradient 2 values in DMN regions and QA values of fibers including the AF, IFOF, AC, corpus callosum, and cingulum (Figure 4A). No significant correlations were identified for Cont_PFCl_6, the region associated with line bisection deviation. To confirm the robustness of these associations, partial correlation analyses were performed between QA and functional gradient values, adjusting for lesion volume and the presence of visual field deficits. These analyses confirmed the results: Cont_PFCl_2: r = 0.568, p = 0.005, DMN mean: r = 0.544, p = 0.007 (Figure 4B)

**Figure 4.**
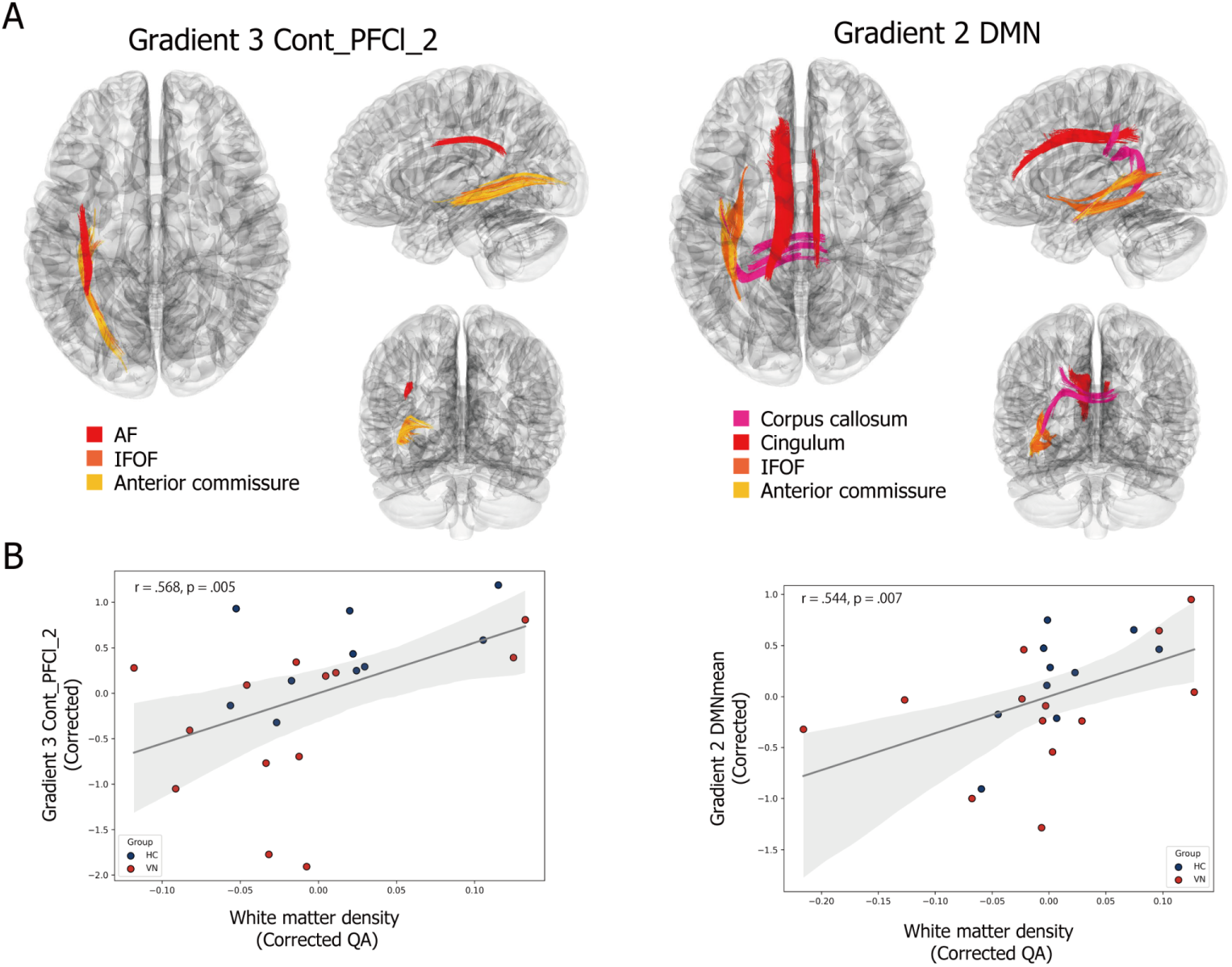
Changes in dispersion along the gradients are associated with underlying white matter structural integrity in both controls and patients. (A) White matter tracts linked to functional gradient alterations. Left: WM tracts positively correlated with Cont_PFCl_2, which showed group-level differences along Gradient 3. Identified WM tracts include the arcuate fasciculus (AF), inferior fronto-occipital fasciculus (IFOF), and anterior commissure. Right: WM tracts correlated with the default mode network (DMN), which were related to neglect severity along Gradient 2. Significant WM tracts include the corpus callosum, cingulum, IFOF, and anterior commissure. (B) Partial correlations between white matter density (corrected QA values) and gradient scores, across both HC and neglect. Gradient scores and white matter density were corrected for nuisance variables, including lesion volume and the presence of visual deficits.

### Individual analyses of functional gradients

Table 2 summarizes the differences with 95% confidence intervals between each neglect patient and the HC group in the ROI of the left lateral PFC, where a significant group difference was observed. For 11 of the 13 patients, the 95% confidence intervals did not include zero, thus showing that these patients followed the general pattern observed in the neglect group.

**Table 2.** The degree of deviation from the HC group in the left lateral prefrontal cortex for each neglect patient.

| ID | Estimate | 95%CI_low | 95%CI_high |  |
| --- | --- | --- | --- | --- |
| Patient 1 | -3.702 | -5.471 | -1.896 | * |
| Patient 2 | 0.003 | -0.616 | 0.621 |  |
| Patient 3 | -0.984 | -1.729 | -0.202 | * |
| Patient 4 | -0.552 | -1.209 | 0.130 |  |
| Patient 5 | -0.784 | -1.488 | -0.057 | * |
| Patient 6 | -0.971 | -1.715 | -0.192 | * |
| Patient 7 | -1.900 | -2.947 | -0.823 | * |
| Patient 8 | -2.511 | -3.795 | -1.198 | * |
| Patient 9 | -4.579 | -6.736 | -2.401 | * |
| Patient 10 | -0.796 | -1.502 | -0.058 | * |
| Patient 11 | -3.195 | -4.754 | -1.617 | * |
| Patient 12 | -4.856 | -7.131 | -2.565 | * |
| Patient 13 | -0.897 | -1.619 | -0.138 | * |

\* 95% confidence intervals not crossing zero.

## Discussion

In this resting-state fMRI study of patients with chronic left spatial neglect, we used a novel functional gradient analysis to investigate macroscale network reorganization, focusing exclusively on the non-lesioned left hemisphere. We deliberately excluded the lesioned hemisphere due to the extensive and heterogeneous anatomical damage present in the majority of patients, which can affect signal quality and fMRI analyses. This approach allowed us to examine indirect, diaschisis-like effects of the lesion on left-hemisphere connectome, although it precludes direct assessment of interhemispheric dynamics. Our study aimed to describe associations between functional gradients, behavioral performance, and structural connectivity. We identified altered functional segregation within key control and default mode networks, as well as structural-functional coupling involving the inferior fronto-occipital fasciculus. Our results suggest that alterations in the balance between compensatory and maladaptive mechanisms in the left hemisphere may be related to persistent neglect behaviors.

While these observations are exploratory, they provide insights into how left-hemisphere network organization could influence attentional performance after right-hemisphere stroke and may inform future investigations into targeted therapeutic strategies, such as neuromodulation. Reduced functional segregation in the left lateral prefrontal cortex, a key node of the frontoparietal control network, was associated with persistent greater rightward deviation in line bisection. This observation extends and specifies convergent evidence from magnetoencephalography (Rastelli et al., 2013), lesion (Vuilleumier et al., 1996) and neuromodulation studies (Oliveri et al., 1999; Zhao et al., 2025), implicating the left PFC in the pathophysiology of left neglect. In particular, Zhao et al., (2025) showed that facilitatory theta burst stimulation of the left PFC improved neglect and enhanced inter-hemispheric connectivity, suggesting a possible mechanism linked to inter-hemispheric communication. These findings suggest that the left PFC may play a dual role, potentially supporting compensatory processes or, conversely, reinforcing maladaptive hemispheric asymmetry, depending on broader network dynamics (Bartolomeo, 2019, 2021). Notably, associations were strongest for line bisection, which may reflect the sensitivity of this task to spatial perceptual bias in chronic neglect, whereas other measures did not show significant correlations.

Beyond its traditional association with language, the left PFC is a domain-general hub engaged in working memory, arousal, consciousness, and visual imagery (Bor & Owen, 2007; Del Cul et al., 2009; Dijkstra et al., 2025). Direct stimulation of this region can elicit hallucinations (Blanke et al., 2000), and neuropsychological evidence in somatoparaphrenic patients suggests that its functional dysconnection with impaired right-hemisphere networks may promote delusional misattributions of self-body (Bartolomeo et al., 2017; Halligan et al., 1995; Saetta et al., 2021). The present evidence indicates that the clinical impact of left PFC activity depends on the balance of integration and segregation within large-scale attention and control networks. On the one hand, insufficient segregation in the left PFC may diminish its ability to flexibly redistribute attentional resources. On the other hand, inappropriate hyperintegration may also be maladaptative. These disruptions of the integration–segregation balance may cause left-hemisphere perceptual decision systems (Heekeren et al., 2004) to receive misleading information, resulting in a lack of or inappropriate responses to left-sided events (Bartolomeo et al., 2025). Therapeutic modulation may thus be most effective when aimed at normalizing network balance rather than simply increasing or decreasing left PFC activity (Bartolomeo, 2019).

Additionally, we found that the pathologically increased segregation of the DMN correlated with rightward attentional bias in line bisection. Given that DMN activity is typically suppressed during externally oriented, attention-demanding tasks (Broyd et al., 2009; Menon, 2023), excessive segregation of this network may reflect an intrusion of internally oriented processes (mind-wandering, self-referential mentation) that compete with external attentional control. This interpretation is supported by evidence linking DMN hyperactivity to lapses in sustained attention (Bonnelle et al., 2011; Weissman et al., 2006) and by primate stroke models in which abnormally enhanced DMN segregation is related to poorer recovery (Nashed et al., 2024). Thus, our results suggest that imbalanced DMN dynamics may limit behavioural compensatory mechanisms in chronic neglect (Bartolomeo, 1997; Takamura et al., 2016; Tham et al., 2000) by interfering with attention to the external environment, although further work is required to determine whether this alteration is specific to neglect or reflects stroke severity more broadly.

Our results specify independent lines of evidence in explaining the striking hemispheric asymmetry of neglect—an unresolved question in clinical neuroscience for nearly a century (Brain, 1941). In the healthy brain, converging evidence from resting-state connectivity (Gotts et al., 2013) and network controllability analyses (Ben Messaoud et al., 2025) indicates a fundamental hemispheric asymmetry in large-scale brain organization. The left hemisphere shows a bias toward communicating within itself, supporting language and motor functions through predominantly segregated intrahemispheric interactions. In contrast, the right hemisphere exhibits a more integrative profile, engaging both intra- and interhemispheric connections—particularly with the left hemisphere. This fundamental asymmetry provides a mechanistic basis for the disproportionate impact of right-hemisphere damage on spatial and attentional functions. It further specifies the recent anatomofunctional model according to which conscious perception depends on the right hemisphere’s capacity to prioritize sensory information and transmit it to left-hemisphere decision systems (Bartolomeo et al., 2025). In this view, right-hemisphere damage critically disrupts both local processing and the interhemispheric information flow necessary for awareness, thereby explaining its disproportionate impact on neglect and conscious access. The present evidence refines this model by stressing the pivotal role of the left prefrontal cortex.

In addition to the SLF, damage to a more ventral caudorostral pathway—the right-hemisphere inferior fronto-occipital fasciculus (IFOF)—has also been associated with left spatial neglect (Urbanski et al., 2008). Our structural-functional coupling analysis suggested a dual role of its left-hemisphere homolog in neglect pathophysiology. Higher axonal density of the left-hemisphere IFOF was positively associated with gradient values in the left PFC (Gradient 3) and DMN (Gradient 2). This suggests that preserved structural connectivity may simultaneously contribute to compensatory reorganization within control networks while also supporting maladaptative segregation of the DMN. Similar dual contributions have been reported in motor recovery and aphasia, where the controlesional hemisphere provides both beneficial redundancy and in some cases, interference (Kaufmann et al., 2024; Saur et al., 2006). Whether the observed correlations reflect premorbid IFOF architecture or post-stroke structural remodeling cannot be resolved in our cross-sectional design. Emerging microstructural imaging techniques and longitudinal research may clarify whether pathological changes in association tracts precede or follow functional reorganization.

In addition to the involvement of intrahemispheric pathways, previous studies underscore the critical role of the corpus callosum, the brain’s largest commissural fiber bundle. The integrity of the corpus callosum facilitates interhemispheric communication, which is essential for recovery of severe forms of spatial neglect (Kaufmann et al., 2024; Lunven et al., 2015, 2019). Damage to the corpus callosum—particularly in its posterior segments connecting parietal regions—is associated with poorer recovery and the persistence of neglect (Lunven et al., 2015), presumably because it disrupts interhemispheric interactions that support adaptive reorganization. Moreover, frontal callosal fibers may support adaptive compensatory mechanisms through communication between motor systems after prism adaptation, a visuomotor rehabilitation technique (Bartolomeo et al., 2017). Our study focused on the intact left hemisphere, thus precluding direct assessment of interhemispheric functional connectivity. However, the coupling observed between the left IFOF and DMN gradients may interact with callosal integrity to influence the degree of functional segregation or integration across hemispheres. Future studies incorporating within- and between-hemisphere structural connectivity, including detailed corpus callosum mapping, are warranted to disentangle these relationships and better contextualize compensatory versus maladaptive mechanisms underlying neglect recovery.

Previous studies have demonstrated that changes in functional gradients are associated with disease-specific alterations in brain network architecture across neurological, psychiatric, and neurodevelopmental disorders (Dong et al., 2023; Hong et al., 2019; Rosa et al., 2024; Tan et al., 2023). In particular, stroke leads to marked changes in functional connectivity gradients, especially affecting those that emphasize specific networks, rather than the networks themselves (Bayrak et al., 2019). This aligns with our findings, in which gradients 2 and 3 were associated with the presence and severity of neglect.

Our results highlight that plasticity after neurological disease is not limited to spontaneous reorganization of spared tissue. Rather, it reflects the interplay of residual structural pathways, compensatory recruitment, and potentially maladaptive network hyperactivity. Specifically, the left prefrontal cortex may function not only as a compensatory resource but also as a pathological reinforcer of hemispheric asymmetry, depending on the underlying connectivity dynamics. Recognizing this dual potential is critical for developing interventions. In light of recent theoretical proposals (Bartolomeo, 2019) and neuromodulation findings (Zhao et al., 2025), strategies aimed at normalizing—rather than simply enhancing or inhibiting—left-hemisphere activity and inter-hemispheric connectivity may prove more effective.

Some limitations should be noted. First, our sample size is relatively small and lacks a control group of right hemisphere-lesioned patients without neglect limiting claims about specificity. However, individual analyses confirmed the general pattern in 11 of the 13 examined patients. Second, only line bisection scores were related to the functional gradient. Third, lesion size posed challenges for fMRI preprocessing, partially mitigated by focusing analyses to the intact hemisphere. While this restriction excludes interhemispheric dynamics, it also provides a novel view of reorganization in severely affected patients, who are often underrepresented in imaging studies. To advance this line of work, future research should include longitudinal designs spanning the subacute to chronic phases of recovery (Ramsey et al., 2016; Siegel et al., 2018), matched lesion controls without neglect, and integration with advanced neurophysiology (Chea et al., 2024; Massimini et al., 2024). Such approaches will help disentangle compensatory from maladaptive mechanisms and may guide precision therapies tailored to individual connectivity profiles.

In conclusion, our study demonstrates that alterations in left-hemisphere network organization, particularly within the non lesioned left lateral prefrontal cortex and default mode networks, are associated with rightward attentional bias in patients with chronic left neglect. Reduced functional segregation in the left prefrontal cortex as part of the frontoparietal control network and increased segregation within the DMN were both linked to greater rightward attentional bias, potentially reflecting a combination of maladaptive compensatory processes. Structural integrity of the left IFOF may modulate these network patterns, supporting either compensatory or maladaptive connectivity depending on the broader functional context. Overall, these findings provide a framework for future studies aimed at disentangling compensatory and maladaptive mechanisms and may inform targeted interventions to optimize network-level function after stroke.

## CRediT authorship contribution statement

**YT**: Conceptualization, Methodology, Software,Validation, Formal analysis, Data Curation,Writing - Original Draft, Writing - Review & Editing, Visualization; **ML**: Conceptualization, Methodology, Investigation, Resources, Data Curation, Writing - Review & Editing, Supervision; **YS**: Software, Writing - Review & Editing; **CB**: Investigation, Resources; **SOC**: Conceptualization, Methodology, Validation, Writing - Review & Editing, Supervision; **PB**: Conceptualization, Resources, Writing - Review & Editing, Supervision, Project administration.

## Acknowledgments

During the preparation of this work, the authors used ChatGPT4 for proofreading and coding. After using this tool, the authors reviewed and edited the content. The authors take full responsibility for the content of the publication.

## Data availability

All the anonymised materials are available on the open science framework (https://osf.io/ztdy4).

